# A critical assessment of LGN segmentation quality based on volume, contour, and location

**DOI:** 10.64898/2026.10.19.759668

**Authors:** Jan-Luca Schröder, Omer Faruk Gulban, Christa Müller-Axt, Rainer Goebel, Jan W. Kurzawski

**Affiliations:** Faculty of Psychology and Neuroscience, Maastricht University, The Netherlands; Faculty of Psychology, Dresden University of Technology, Dresden, Germany; Max Planck Institute for Human Cognitive and Brain Sciences, Leipzig, Germany; Brain Innovation, Maastricht, The Netherlands

**Keywords:** Lateral geniculate nucleus (LGN) -Thalamus -Segmentation -MRI -Evaluation -Subcortex -Anatomy, Brain

## Abstract

Reliable segmentation of the lateral geniculate nucleus (LGN), a small thalamic relay nucleus essential for visual processing, is necessary for characterizing its anatomy in healthy populations and structural alterations in neurological disorders. However, accurate delineation remains challenging on routinely acquired T1-weighted images because the LGN is small, deeply located, and has limited contrast relative to surrounding thalamic tissue. Although manual segmentation remains the reference standard, it is time consuming and susceptible to rater variability, limiting its use in large datasets. Automated approaches may therefore provide a more scalable alternative. In this study, we performed a systematic benchmarking of seven automated approaches: FreeSurfer, NextBrain, DeepThalamus, HIPS-THOMAS, MASSP 2.0, BigBrain alignment, and manual novice segmentation against expert-defined ground truth. We applied a multi-metric evaluation incorporating the Dice similarity coefficient, Hausdorff distance and balanced average Hausdorff distance, volumetric similarity, and Euclidean distance. Together, these measures evaluate overlap, contour, volume, and location against expert manual segmentations. Our results indicate that HIPS-THOMAS and DeepThalamus outperformed the other automated methods in all metrics. However, manual segmentation retained high precision despite exhibiting greater inter-subject variance than the top-performing automated segmentation tools. The different metrics do not necessarily report the same order of quality, highlighting the need of using multiple metrics for evaluation segmentation quality.

## Introduction

The human lateral geniculate nucleus (LGN) is the principal thalamic relay of the visual pathway (Sherman & Guillery, 1996; Van Horn et al., 2000). Beyond relaying retinal input to primary visual cortex, the LGN receives extensive corticothalamic and thalamic feedback (Briggs, 2020; Ghodrati et al., 2017) and contributes to visual processing (Casanova & Chalupa, 2023; Gamal & Eldawlatly, 2025; Ghodrati et al., 2017), attention (Ling et al., 2015; Poltoratski et al., 2016; Shah et al., 2022), and sensory modulation (Alitto et al., 2026; Kastner et al., 2006; Shah et al., 2022). Recent advances in ultra-high-field MRI have enabled increasingly detailed in vivo investigations of the human LGN, facilitating both anatomical mapping and functional studies of this small but complex thalamic nucleus (Denison et al., 2014; Jia et al., 2023; Keuken et al., 2018; Müller-Axt et al., 2025).

The human LGN exhibits substantial inter-individual variability in volume, shape, and location (Andrews et al., 1997; Halpern et al., 1999; Liu et al., 2026). Capturing this variability is important for studies of both the anatomy and function of the visual system and therefore requires reliable subject-specific segmentation. Segmentation inaccuracies may obscure genuine anatomical differences or introduce measurement bias. Detailed characterization of the LGN is also clinically important, because abnormalities of this nucleus have been associated with neurological and ophthalmological disorders, including glaucoma (Chen et al., 2025; Kosior-Jarecka et al., 2020; Yang et al., 2025), idiopathic blepharospasm (Meoni et al., 2024), Alzheimer’s disease (Andrews et al., 1997; Halpern et al., 1999; Liu et al., 2026), and multiple sclerosis (Korsholm et al., 2007; Papadopoulou et al., 2019). Moreover, the first successful recordings of human LGN cells were recently reported (Self et al., 2025), paving the way for a visual prosthesis targeting the LGN (van Hoof et al., 2025).

Collectively, these considerations underscore the functional and clinical importance of the LGN and the need for its accurate identification is essential in both basic neuroscience and clinical research. Yet its small size, central location, and anatomical complexity make reliable segmentation challenging (Saranathan et al., 2015), creating a need for robust and reproducible methods.

### Segmentation of the LGN

Automated segmentation performance depends strongly on the MRI contrast. Most methods have been developed for T1-weighted structural images, anatomical data most commonly acquired in both research and clinical practice. However, the LGN has little contrast relative to surrounding thalamic tissue in T1-weighted images, making accurate segmentation challenging. Additional MRI contrasts, including T2*-weighted or diffusion-weighted (Kitajima et al., 2015; Schmidt et al., 2018; Wang et al., 2015; Y. Zhang et al., 2012) and proton-density-weighted images (Aldusary et al., 2019; Devlin et al., 2006; Welbourne et al., 2025), can improve delineation but require additional scan time and may provide limited spatial resolution (Behrens et al., 2003; DeSimone & Schneider, 2019; Giraldo-Chica & Schneider, 2018). Moreover, these contrasts are rarely included in existing, publicly available datasets, limiting the applicability of methods that rely on them. We therefore focused on T1-weighted MR images to evaluate methods with broad applicability to existing and routinely acquired data.

Current approaches to LGN segmentation can be broadly divided into three categories: First, manual segmentations done by an expert segmenter, which provide highly individualized segmentations tailored to each subject’s anatomy. These are seen as the gold standard (Akudjedu et al., 2018; Müller-Axt et al., 2021; Welbourne et al., 2025). However, manual segmentation is time-consuming, requires considerable anatomical expertise, and remains susceptible to intra-and inter-rater variability.

Second, some methods rely solely on spatial normalization, transferring an LGN label from a population template or histological atlas into individual subject space. Examples include MNI-space atlases like the one in Krauth et al. (2010) and the histology-based BigBrain data (Amunts et al., 2013). Although computationally simple, these approaches depend heavily on accurate image registration, often differ in exact terminology (Mai & Majtanik, 2019), and may not adequately capture individual anatomical variability.

Third, several automated methods have been developed to deal with large datasets where manual segmentation is not feasible. These include probabilistic atlas-based approaches such as FreeSurfer’s Bayesian thalamic segmentation (Iglesias et al., 2018), (multi-)atlas methods such as HIPS-THOMAS (Vidal et al., 2024) or MASSP 2.0 (Bazin et al., 2025), and more recent deep learning approaches including DeepThalamus (Ruiz-Perez et al., 2024), THOMAS-CNN (Williams et al., 2022), and NextBrain (Casamitjana et al., 2025). While these methods all substantially reduce manual effort, they differ in their underlying data sources, specific approaches, required image contrasts, and expected segmentation accuracy.

The growing number of available methods makes selecting an appropriate LGN segmentation approach increasingly difficult. Post-mortem measurements provide anatomical reference data but may be affected by differences between in vivo and post-mortem tissue and are inherently limited to anatomical measurements. Expert manual segmentations therefore provide the closest available approximation to an in vivo ground truth, although its performance must still be interpreted carefully. Regarding the (semi-)automated approaches, most studies on evaluating the proposed methods report favorable results, but their validation on different datasets, with different reference standards and evaluation metrics prevents direct comparison and offers limited guidance for method selection. Moreover, evaluations commonly limit assessment of segmentation quality to a single metric (Müller et al., 2022; Taha & Hanbury, 2015), most often the Dice coefficient (Popovic et al., 2007; Zhang et al., 2025). Dice quantifies spatial overlap but does not distinguish errors in contour, location, or volume (Taha & Hanbury, 2015). Segmentation quality is therefore multidimensional: two segmentations can have similar Dice scores yet differ meaningfully in boundary accuracy, anatomical position, or volumetric bias. These differences are particularly consequential when studying individual variability or clinical effects in a small structure such as the LGN.

A comprehensive comparison of current LGN segmentation methods should assess complementary dimensions of performance rather than rely on a single metric. Fig. 1 outlines the conceptual framework of this study, including the challenges of LGN segmentation, the segmentation approaches evaluated, the dimensions of segmentation quality, and the practical effort required. In this study, we evaluate seven current subcortical segmentation methods using high resolution 7T T1-weighted MR images and compare their LGN segmentations with expert manual segmentations. We also examine whether greater practical effort is associated with higher segmentation quality. By jointly quantifying overlap, contour, volume, and location, we aim to identify method-specific strengths and limitations and provide practical guidance for selecting an LGN segmentation approach for T1-weighted MR images.

**Fig 1.**
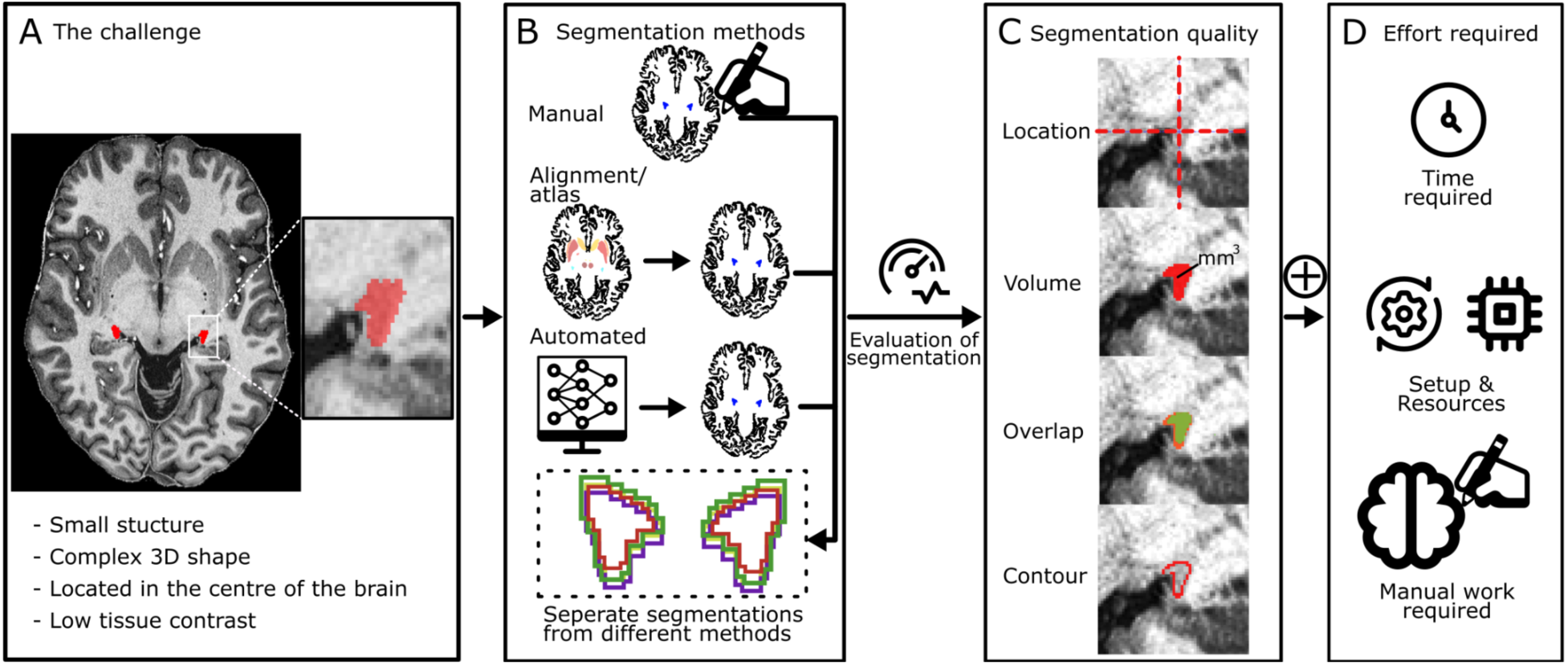
Conceptual framework for evaluating LGN segmentation. **A:** A description of the challenge of segmenting the human LGN. **B:** Example of approaches of segmentation methods. **C:** Quallity is defined as a segmentation being highly accurate and precise in terms of contour, volume, and location. D: Effort is defined here as the cumulative time required to generate a segmentation. This includes the active manual labor involved in the segmentation process, the time allocated for method configuration, and the operational requirements necessary for execution

## Methods

### MRI data

We used a dataset of 28 healthy subjects, comprising 3D-MP2RAGE acquisitions including T1-weighted uniform (UNI) images (mean age 25.9 ± 3.8 years, 15 female) and quantitative T1 maps. The data were acquired on a MAGNETOM 7T MRI system (Siemens Healthineers, Forchheim, Germany) with a 24-channel Nova head coil (NOVA Medical, Wilmington MA, USA) with a 0.5 mm isotropic resolution (Tardif et al., 2016). One participant was excluded due to a later diagnosis of developmental dyslexia. All images were upsampled at a resolution of 0.4 mm isotropic using a rigid (6-parameter) transformation (Müller-Axt et al., 2021; Tardif et al., 2016). The images were preprocessed using the steps described in (Tardif et al., 2016). This dataset was used for its high spatial resolution, which makes capturing small changes in the LGN boundaries possible.

### Segmentation

We applied several state-of-the-art (semi-)automated segmentation methods, as well as an alignment-based method, and manual segmentation onto the data. Based on these segmentations, we calculated descriptive metrics like the volume, surface area and sphericity, the latter as defined by (Wadell, 1935), of each segmentation. In the following, we are going to describe the processing steps taken with each segmentation method. For each segmentation method, the LGN segmentations were saved individually per hemisphere.

#### FreeSurfer

The skull-stripped MRI data were preprocessed with FreeSurfer (version 8.2.0) using *recon-all* with the *no-skullstrip* flag. The output of this was then processed using *segmentThalamicNuclei.sh*, as described in (Iglesias et al., 2018). Here, the raw unaltered data was read into the script as a NIfTI file, in addition to the subject folder processed with *recon-all*. Due to the limited resolution of the *recon-all* subject data, the segmentation methods native resolution, the output of the segmentation was at a resolution of 0.5 mm isotropic. This output was upsampled to 0.4 mm isotropic using *Nearest Neighbor* interpolation and then resliced into the subject space. The segmentations of the LGN were then converted from .*mgz* into .*nii.gz* images using *mri_convert*.

#### NextBrain

The raw T1w images were processed as NIfTI images using the *mri_histo_atlas_segment_fireants* function, bundled with FreeSurfer (version 8.2.0), as described in (Casamitjana et al., 2025). The LGN segmentations were then isolated and resliced into the subject’s image space using *Nearest Neighbour* interpolation.

#### DeepThalamus

The subject’s data were processed using the DeepThalamus pipeline (Ruiz-Perez et al., 2024) through the online tool volBrain (Manjón & Coupé, 2016). Here, the raw images were uploaded and processed individually. The processed segmentations were then downloaded from the volBrain server.

#### HIPS-THOMAS

The individual T1-weighted images were processed using the HIPS-THOMAS segmentation pipeline described in (Vidal et al., 2024) using the provided Docker container and then extracted as binary segmentations.

*MASSP 2.0*

LGN segmentation was performed using MASSP 2.0 (Bazin et al., 2025), implemented in the Nighres toolbox (Huntenburg et al., 2018). Quantitative T1 maps were converted to R1 (=1/T1) maps. A shared registration transformation was estimated using two image pairs: the AHEAD R1 template and each participant’s R1 image, and the MNI09b T1-weighted template and each participant’s MP2RAGE UNI image. The MNI09b template was resampled onto the AHEAD R1 grid before registration. The resulting mapping was supplied to MASSP 2.0, with the participant’s R1 image as the sole intensity contrast for segmentation.

#### BigBrain

For a strictly alignment-based segmentation approach, we manually segmented the LGNs on the MNI-aligned BigBrain data at a resolution of 100 *μm* isotropic (Amunts et al., 2013; Xiao et al., 2019) by a single segmenter due to the clear contrast of the LGN to the surrounding tissue. The BigBrain image was then downsampled to 400 *μm* isotropic resolution. The individual anatomy was then aligned to the MNI template (ICBM152 2009b symmetric) using ANTs with non-linear SyN normalization (Tustison et al., 2021). We then transformed the manual LGN segmentations onto the subject’s native space using the inverse-transformation using *Nearest Neighbour* interpolation.

#### Manual Segmentations

Lastly, manual segmentations in the subject’s native space (manual novice segmentation) by a single segmenter. For this, the individual images were loaded into ITK-SNAP (*version 4.4.0*) (Yushkevich et al., 2006), the contrast was adjusted to make the visual detection of the boundaries of the LGN easier, and then manually drawn on a slice by slice level using a drawing tablet. Importantly, the segmenter had limited prior experience in segmenting the LGN. Therefore, we classify this here as a ’novice’ segmentation, in order to contrast it to the expert segmentation used as the ground truth.

### Segmentation evaluation

To evaluate the performance of the various segmentation methods against the expert-defined ground truth (Müller-Axt et al., 2021), we utilized the EvaluateSegmentation software (Taha & Hanbury, 2015). Five complementary metrics were used to capture different aspects of segmentation accuracy: the Hausdorff distance (HD), the balanced average Hausdorff distance (baHD) (Aydin et al., 2021), volumetric similarity, the Euclidean distance between the centroids (centers of mass), and the Dice similarity coefficient. The Hausdorff distance and balanced average Hausdorff distance were utilized both to characterize different aspects of the contour. The Hausdorff distance is highly sensitive to outliers, making it ideal for detecting the maximum local error, whereas the balanced average Hausdorff distance offers a more robust measure of the average distance between the contours. Volumetric similarity was included to evaluate the consistency of the total volume regardless of spatial overlap.

The volumetric similarity is defined as 1 minus the volumetric difference divided by the sum of the volumes (Taha & Hanbury, 2015). To quantify the global positional shift between the segmentations, we calculated the Euclidean distance between the centroids. This provides a measure of positional accuracy that is independent of contour fluctuations. Finally, the Dice coefficient was used to quantify the spatial overlap between the segmentations. The definition of the metrics can be found in (Taha & Hanbury, 2015). We decided to incorporate this selection of metrics, as individual metrics do not cover all aspects of a segmentation and might miss specific aspects relevant for a specific application.

### Statistical analysis

To evaluate if there is an effect of the segmentation method on the different metrics, we performed separate non-parametric Friedman tests.

## Results

Given the increasing number of automated LGN segmentation methods, we systematically evaluated their performance relative to expert manual segmentations (Müller-Axt et al., 2021). Different segmentation methods produce very different estimates of the anatomy of the LGN, even before comparing them to a ground truth. As a first step, we examined descriptive statistics for each LGN segmentation method to characterize variation in key anatomical features and to perform comparison with previous LGN segmentation studies.

### Descriptive statistics

The expert drawings of the LGN exhibit a median volume of 120.74 mm^3^ (L: 120.38 mm^3^ and R: 121.09 mm^3^), a median surface area of 176.81 mm^2^ (L: 176.54 mm^2^; R: 177.14 mm^2^) and a sphericity of 0.66 (L: 0.65; R: 0.66). The manual novice and BigBrain segmentations, followed by the HIPS-THOMAS segmentation reported the closest volume to the manual expert segmentation, while the other methods reported either larger or smaller volumes, with MASSP 2.0 reporting the smallest and FreeSurfer the largest volumes (Table 1). For the surface area, the manual novice segmentation, MASSP 2.0 and NextBrain reported the most similar values to the expert manual segmentation, while FreeSurfer reported the largest and DeepThalamus the smallest surface area. For the sphericity, the methods differ strongly between each other. FreeSurfer and HIPS-THOMAS report the closest sphericity values to the expert manual segmentations. DeepThalamus reports the most spherical segmentation and NextBrain and MASSP 2.0 the least spherical segmentation. Moreover, as these statistics also differ within each method between the left and right side, we further investigated the volumetric similarity (see Fig. S2). These results show us how the different segmentations differ in terms of basic descriptive statistics. However, this does not provide an indication of the quality of the resulting segmentation. Therefore we next evaluated how each method compared to the expert-defined ground truth in terms of overlap, contour, volume, and location.

**Table 1.**
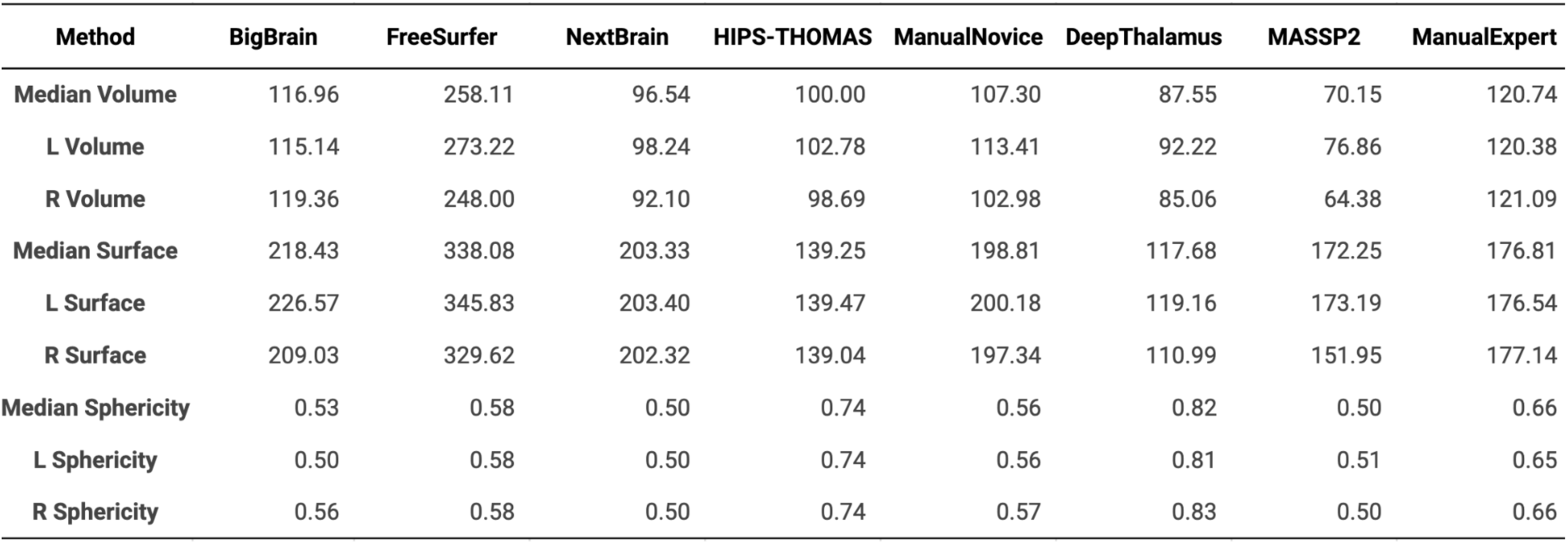
Descriptive metrics of the segmentation methods. Descriptive metrics, i.e. the median volume (in mm^3^), the surface area (in mm^2^) and the sphericity (1 = spherical, 0 = non spherical) for the left and right LGN for the different segmentation methods

| Method | BigBrain | FreeSurfer | NextBrain | HIPS-THOMAS | ManualNovice | DeepThalamus | MASSP2 | ManualExpert |
| --- | --- | --- | --- | --- | --- | --- | --- | --- |
| Median Volume | 116.96 | 258.11 | 96.54 | 100.00 | 107.30 | 87.55 | 70.15 | 120.74 |
| L Volume | 115.14 | 273.22 | 98.24 | 102.78 | 113.41 | 92.22 | 76.86 | 120.38 |
| R Volume | 119.36 | 248.00 | 92.10 | 98.69 | 102.98 | 85.06 | 64.38 | 121.09 |
| Median Surface | 218.43 | 338.08 | 203.33 | 139.25 | 198.81 | 117.68 | 172.25 | 176.81 |
| L Surface | 226.57 | 345.83 | 203.40 | 139.47 | 200.18 | 119.16 | 173.19 | 176.54 |
| R Surface | 209.03 | 329.62 | 202.32 | 139.04 | 197.34 | 110.99 | 151.95 | 177.14 |
| Median Sphericity | 0.53 | 0.58 | 0.50 | 0.74 | 0.56 | 0.82 | 0.50 | 0.66 |
| L Sphericity | 0.50 | 0.58 | 0.50 | 0.74 | 0.56 | 0.81 | 0.51 | 0.65 |
| R Sphericity | 0.56 | 0.58 | 0.50 | 0.74 | 0.57 | 0.83 | 0.50 | 0.66 |

### Analysis of the segmentation methods

To evaluate the performance of the segmentation methods, five complementary metrics were chosen to quantify different aspects of segmentation quality. Some example segmentations can be seen in Fig. 3. The influence of the segmentation method on the performance in each metric was evaluated using the Friedman test, which indicated significant main effects across all five metrics (Hausdorff Distance: *Fr(6)* = 117.0, *W* = 0.75, *p* = 6.55e^-23^, Balanced-average Hausdorff Distance: *Fr(6)* = 124.99, *W* = 0.80, *p* = 1.45e^-24^, Volumetric similarity: *Fr(6)* = 114.05, *W* = 0.73, *p* = 2.89e^-22^, Dice coefficient: *Fr(6)* = 129.18, *W* = 0.82, *p =* 1.91e^-25^, Euclidean Distance: *Fr(6)* = 118.04, *W* = 0.76, *p* = 4.21e^-23^).

**Fig 3.**
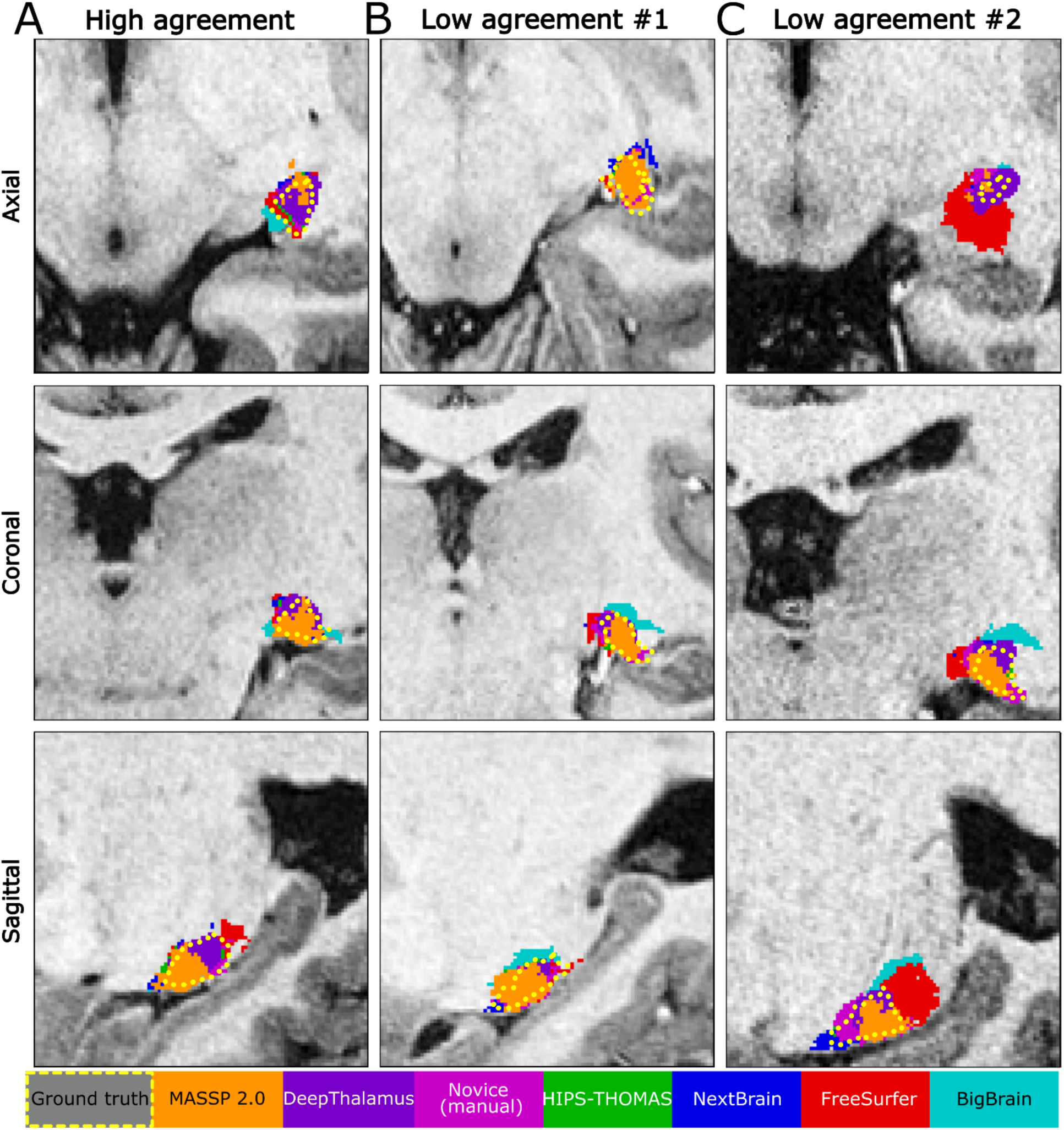
Overview of highest and lowest segmentation agreement between different segmentation methods. **A** displays an axial, coronal and sagittal slice of an example subject where the overall agreement between different segmentation methods is high. **B** shows mislocalization in the axial and sagittal slices. **C** illustrates the effect of contour mismatches and differences in estimated size (and thus volume) between the different segmentation methods. Fig. S1 shows the original T1-weighted images without added segmentations

Regarding boundary and location, the methods followed a consistent ranking. For the Hausdorff distance and the balanced average Hausdorff distance, the order of performance from lowest (best), to highest (worst) value was: manual (novice), HIPS-THOMAS, DeepThalamus, MASSP 2.0, NextBrain, Bigbrain and FreeSurfer. For the Euclidean distance between the centroids, the ranking was slightly shifted, with the manual (novice) segmentation performing best, followed by DeepThalamus, HIPS-THOMAS, MASSP 2.0 and NextBrain, BigBrain, and FreeSurfer.

In terms of volumetric and overlap metrics, the results varied. For volumetric similarity, the highest (best) values were observed for BigBrain, followed by HIPS-THOMAS, NextBrain, manual (novice), DeepThalamus, MASSP 2.0, and FreeSurfer. For the Dice coefficient, the manual (novice) segmentation ranked highest, followed by DeepThalamus, HIPS-THOMAS, MASSP 2.0, NextBrain, FreeSurfer, and BigBrain. Exact values are provided in Table 2 and Fig. 4.

**Fig 4.**
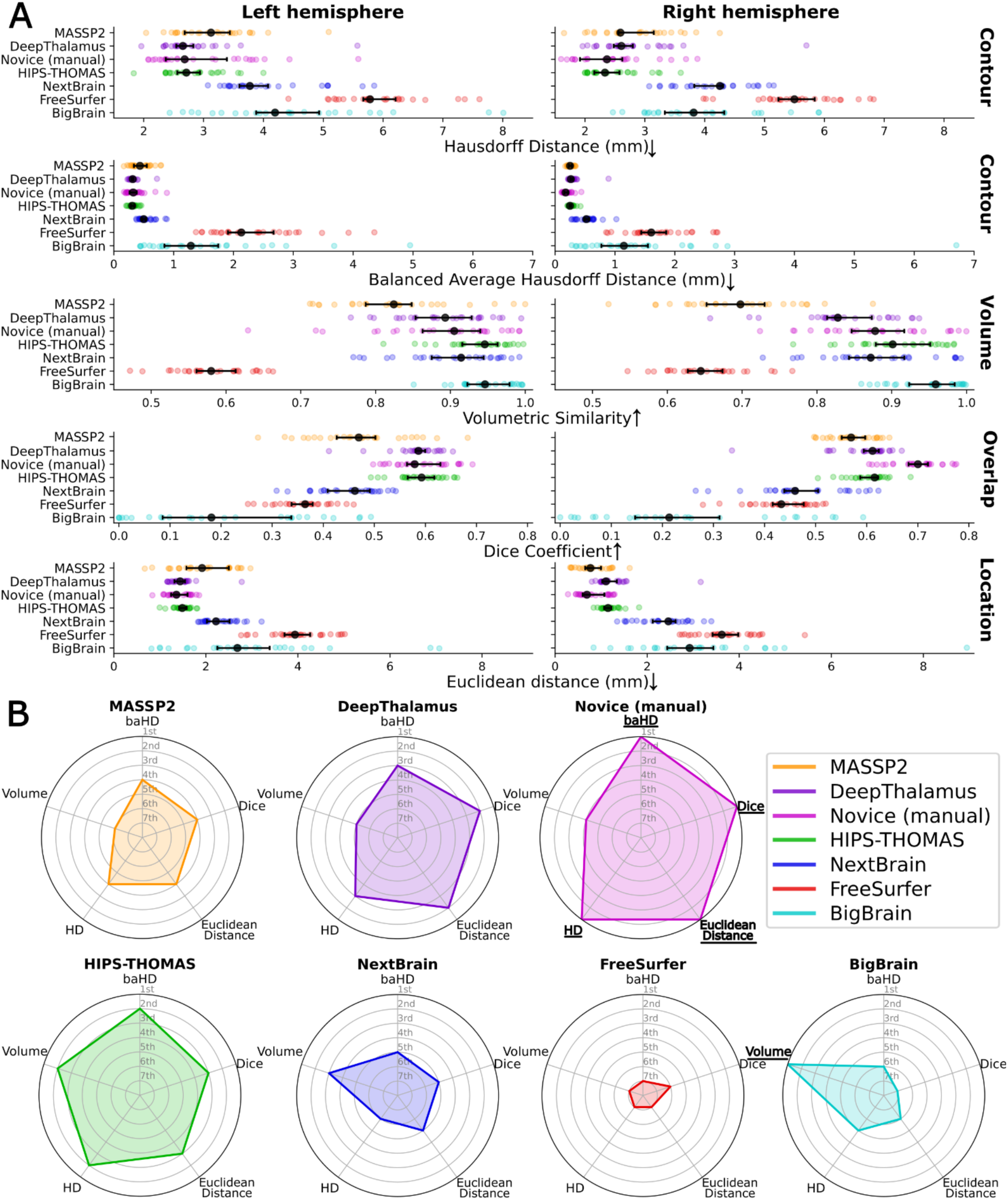
Comparison of different segmentation methods across five metrics, based on the median results for the left and right side LGN at 0.4^3^ mm^3^ image resolution. **A:** The central dot (•) indicates the median value. ↓ indicates lower is better. ↑ indicates higher is better. The error bars indicate the 95% confidence interval. The left column shows the left LGN, the right column the right LGN.**B:** Ranked performance across the different metrics based on the median value of the left and right LGN. Underlined bold metrics indicate that the method performed best in this metric.

**Table 2.** Median values of metrics regarding segmentation quality metrics. The table displays the median values of the segmentation quality metrics regarding each method for the left and right LGN

| Method | BigBrain |  | FreeSurfer |  | NextBrain |  | HIPS-THOMAS |  | Manual (novice) |  | DeepThalamus |  | MASSP 2.0 |  |
| --- | --- | --- | --- | --- | --- | --- | --- | --- | --- | --- | --- | --- | --- | --- |
|  | *L* | *R* | *L* | *R* | *L* | *R* | *L* | *R* | *L* | *R* | *L* | *R* | *L* | *R* |
| Hausdorff Distance (mm) | 4.2 | 3.82 | 5.78 | 5.5 | 3.77 | 4.25 | 2.71 | 2.33 | 2.68 | 2.37 | 2.65 | 2.61 | 3.12 | 2.59 |
| Balanced Average Hausdorff Distance (mm) | 1.29 | 1.15 | 2.13 | 1.61 | 0.5 | 0.52 | 0.31 | 0.25 | 0.32 | 0.18 | 0.32 | 0.26 | 0.44 | 0.25 |
| Volumetric Similarity | 0.95 | 0.96 | 0.58 | 0.64 | 0.91 | 0.87 | 0.95 | 0.9 | 0.9 | 0.88 | 0.89 | 0.83 | 0.82 | 0.7 |
| Dice Coefficient | 0.18 | 0.21 | 0.36 | 0.43 | 0.46 | 0.46 | 0.59 | 0.62 | 0.58 | 0.7 | 0.59 | 0.61 | 0.47 | 0.57 |
| Euclidean Distance (mm) | 2.68 | 2.91 | 3.92 | 3.58 | 2.22 | 2.36 | 1.48 | 1.14 | 1.36 | 0.68 | 1.41 | 1.1 | 1.86 | 0.75 |

## Discussion

Segmenting structures like the LGN is challenging, due to its small size, deep location within the brain and generally due to the limited contrast differences with the surrounding tissue. In this paper we report that segmentation quality is multidimensional. No individual method performed best across all metrics, meaning that the optimal segmentation approach depends on whether priority is given to overlap, contour, volume, or location. HIPS-THOMAS and DeepThalamus performed the best and most consistent across the automated methods, whereas the novice manual segmentation achieved high agreement with the expert-defined ground truth, but exhibited greater subject-level variability. Our findings support a multi-metric evaluation of LGN segmentations, with particular emphasis on the segmentation features most relevant to the scientific or clinical application.

Across the different segmentation methods, LGN volume estimates varied widely, and most approaches fell within the range reported in the literature (Aldusary et al., 2019; Andrews et al., 1997; Lipin et al., 2021; Wang et al., 2015, p. 2015; Welbourne et al., 2025). This range of reported sizes likely reflects either anatomical variability, or effects based on methodological differences in resolution, contrast and in vivo versus post-mortem. Notably, MASSP 2.0, DeepThalamus and NextBrain consistently produced the smallest volumes, whereas FreeSurfer tended to overestimate the size of the LGN. This pattern fits a general tendency of some automated, threshold or atlas-based segmentations to be more conservative regarding the lateral parts of the LGN, while FreeSurfer also seems to include parts of the thalamic tissue surrounding the LGN. The surface area is less clear to interpret, as the LGN has a complex shape with a highly subdivided inner structure (Andrews et al., 1997; Hickey & Guillery, 1979). Therefore, the measured values provide an indirect readout of the overall shape and volume, also linking them to the sphericity findings. Here, most methods yielded a somewhat similar sphericity, however, DeepThalamus and HIPS-THOMAS produced more spherical segmentations than the expert manual segmentations. This suggests that these automated methods lose some of the shape’s complexity, possibly due to some over-regularization. On the other hand, algorithms like MASSP 2.0 and NextBrain produced less spherical segmentations with a larger surface area.

Given the observed hemispheric differences in these morphological characteristics, we further investigated the asymmetry, using the asymmetry index (Fig. S2). The asymmetry index (AI) for the LGN volume is defined as *AI* = (*R* − *L*) / (*R* + *L*). Previous studies reported conflicting results regarding a possible asymmetry of the human LGN, where some studies reported a larger right LGN (Kastner et al., 2004; Li et al., 2012; Müller-Axt et al., 2021; Oishi et al., 2023), while some reported a larger left LGN (Giraldo-Chica et al., 2015; Wang et al., 2015). In our dataset, the manual expert segmentations, as well as the BigBrain segmentation, captured a larger right LGN, while MASSP 2.0, DeepThalamus, NextBrain and FreeSurfer captured a larger left LGN, inverting the direction compared to the ground truth. However, this difference is very small in magnitude, which is why it is plausible that it stems from either resolution of the underlying template or atlas or boundary accuracy difference of the separate methods. HIPS-THOMAS showed no median asymmetry.

The overall differences across descriptive measures underline the limitations of evaluating segmentation quality using a single anatomical feature or metric. Current evaluations often rely primarily on Dice overlap or volume, although other measures, including the balanced average Hausdorff distance, are also used in some studies (Iglehart et al., 2020; Williams et al., 2022, 2024). Considering additional metrics is an important step towards a more comprehensive evaluation; however, each measure captures only specific aspects of segmentation performance when interpreted in isolation. We therefore evaluated how closely each method matched the expert manual segmentations in terms of contour, volume, location, and overlap. This multi-metric assessment supports selection of the most appropriate method according to the specific application and its priority, such as accurate volume estimation, boundary delineation, spatial localization, or overlap.

Applying multi-metric assessment revealed distinct performance profiles across the automated approaches. Across the automated approaches, HIPS-THOMAS and DeepThalamus showed the most favorable overall performance profiles, indicating that they provide the most consistent balance of overlap, contour accuracy, localization, and volume similarity. The comparatively strong performance of HIPS-THOMAS across metrics suggests that it may be a robust choice when a broadly reliable automated LGN segmentation is required. DeepThalamus performed similarly well overall; however, its failure to produce a segmentation for one subject and the extreme result observed in another should be considered when assessing its robustness. NextBrain showed comparatively favorable volume estimates, highlighting that the preferred method may differ when accurate volume estimation is the primary aim.

The novice manual segmentations achieved strong median agreement with the expert manual segmentations across several metrics. Only in terms of volumetric similarity, several other methods achieved values closer to the ground truth. However, the substantially greater subject level variability in some metrics indicates that this agreement was not equally reliable across all participants. This pattern likely reflects case specific challenges, such as low tissue contrast or ambiguous boundaries, which may affect manual delineation differently across individuals and segmenters. In contrast, automated methods apply the same decision rules consistently across participants. These findings illustrate the value of a multi-metric evaluation, as strong median performance in one metric may conceal variability or limitations that become apparent when several complementary measures are considered. Manual segmentation also requires considerable time and expert effort, which further affects its practical suitability for larger datasets.

The alignment-based BigBrain segmentation differed markedly from the other approaches. Its volume estimates closely matched the expert manual segmentations, but its performance was more variable and generally lower for the remaining metrics, particularly Euclidean distance and balanced average Hausdorff distance. This pattern suggests that, in healthy populations without substantial anatomical variation, a simple atlas alignment approach may provide a practical proxy for group level LGN volume estimation. However, it is less suitable for individualized assessment or pathological settings, where local structural changes, such as disease-related atrophy, may not be captured by registration to a general atlas.

We intentionally used whole brain alignment and limited individual refinement to represent a standardized and practically feasible workflow. Similarly, the novice manual segmentations were completed once per subject to provide a consistent comparison across methods. The discrepancy between the accurate volume estimates and poorer spatial performance of the BigBrain segmentation provides a clear example of why segmentation quality cannot be summarized by a single metric. Accordingly, the BigBrain approach may be useful when approximate group-level LGN volume estimates are sufficient, but it should not be assumed to provide accurate individualized contours or spatial localization. The individual data was well aligned in terms of the whole brain, but our results suggest that this is not easily transferable onto how well the alignment worked in terms of the LGN. However, it might suggest a certain relationship between the volume of the LGN and the size of the brain.

In addition to segmentation quality, the effort required to generate each segmentation is an important practical consideration (Fig. 5). Effort is defined here as the cumulative time required to generate a segmentation. This includes the active manual labor involved in the segmentation process, the time allocated for method configuration, and the operational requirements necessary for execution. Quality is defined as a segmentation being highly accurate and precise in terms of contour, volume, and location. Several of the methods chosen require powerful hardware with a large amount of RAM and considerable time to run the segmentation on these high resolution images. This is specifically the case for FreeSurfer, MASSP 2.0 and NextBrain and partially also for HIPS-THOMAS. FreeSurfer also sometimes has difficulties processing very high resolution data (Fortin et al., 2025) due to higher sensitivity to noise, requiring extensive preprocessing to work (Zaretskaya et al., 2018), which is why we pre-processed the data at a lower resolution and loaded the ‘native’ resolution images later back in for the segmentation. Taken together, this limits its applications. HIPS-THOMAS currently requires a x86 processor to work, which could be a limitation for some applications in the future. DeepThalamus on the other hand can be run on the volBrain servers, requiring no specific local hardware. However, due to the online-only nature of the DeepThalamus pipeline, there is no possibility to tweak or adjust processing steps. Furthermore, reliance on external servers excludes sensitive clinical data from local security protocols, reducing its generalizability across all study designs. Non-linear alignment using ANTs (Tustison et al., 2021) of high resolution images, for approaches like the BigBrain alignment, can also require powerful hardware and might take a considerable amount of time.

**Fig 5.**
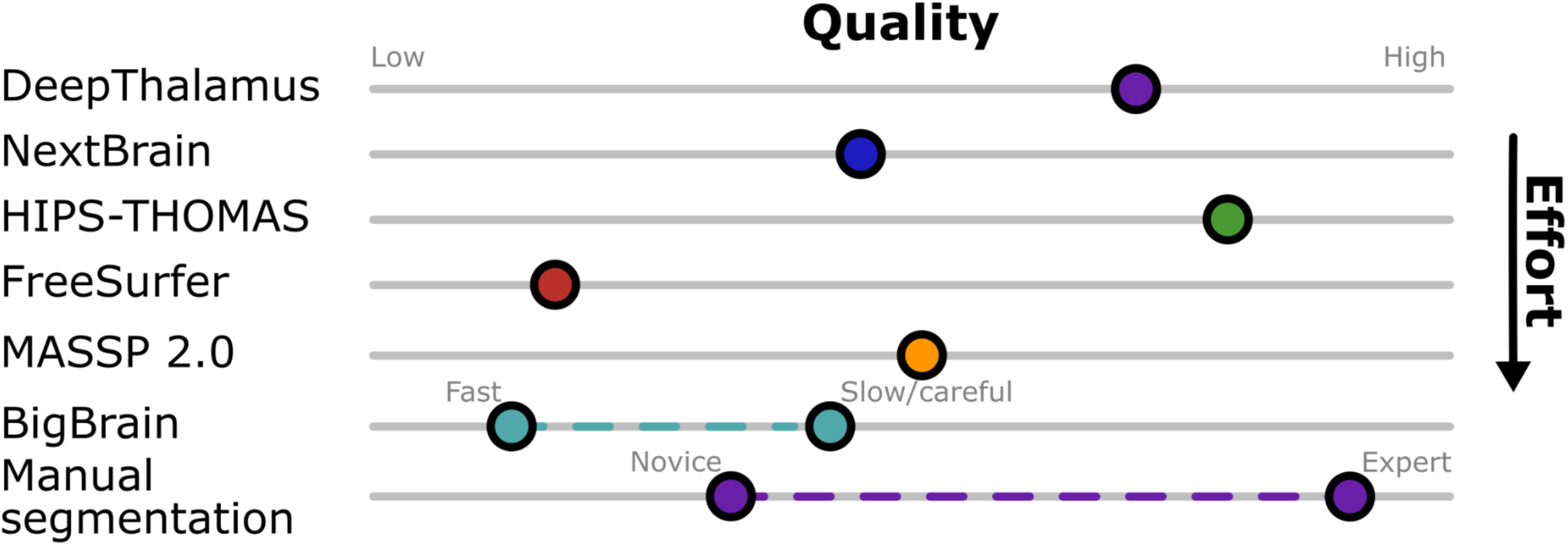
Quality/Effort pay-off. Effort. **Effort** is defined here as the cumulative time required to generate a segmentation. This includes the active manual labor involved in the segmentation process, the time allocated for method configuration, and the operational requirements necessary for execution. **Quality** is defined as a segmentation being highly accurate and precise in terms of contour, volume, and location. The quality score is based on the total derived from summing the rankings of each metric. Two dots for a method indicate a roughly estimated range, dependent on factors indicated. Due to the values being based on rankings, the exact values should be interpreted with some possible variance.

Given these computational considerations, we also assessed whether the high native resolution of the T1-weighted images was necessary for the relative performance of the automated methods. We decided to use high resolution 0.4 mm isotropic data because it helps with manual segmentation and can show LGN contour in more detail. Re-testing the methods on 0.8 mm isotropic resolution (downsampled using ANTs (Tustison et al., 2021) with *BSpline* interpolation) yields the same results in terms of which method performs best in the respective metrics.

Collectively, the data suggest a trade-off between the consistency of automated methods and the anatomical specificity of manual segmentations. The tendency of some automated tools to produce more spherical shapes and symmetric volumes suggests a level of over-generalization, where the algorithms prioritize a ‘standard’ LGN model over the subject-specific irregularities found in the ground truth. However, automated segmentation like HIPS-THOMAS and DeepThalamus create high quality automatic LGN segmentations requiring less effort than individual manual segmentations.

### Limitations

Despite the findings of this study, several limitations must be acknowledged. Due to different datasets having different resolutions and noise levels, the generalizability is naturally somewhat limited. As we used 0.5 mm isotropic data resampled to 0.4 mm isotropic resolution, which is a higher resolution than most collected anatomical images at 7T, our results might not be able to be directly applicable to lower resolution data. However due to current advancements in MRI hardware and sequence development, we believe that within a short timeframe T1-weighted images with a resolution of 0.5 mm isotropic will become more common at high field. Second, we could only evaluate currently available segmentation approaches. While future methodologies may yield superior performance, the multi-metric evaluation framework established here provides a robust baseline and a benchmark against which future automated tools can be compared. Lastly, it is also important to mention that we focused here on the LGN only. Naturally, the performance of the different methods might differ for other structures in the thalamus. Nevertheless, we argue that, using a similar evaluation approach, one could evaluate the segmentation methods for any other structure of interest as well.

## Conclusion

In this study, we investigated how to critically evaluate different segmentation approaches and applied this onto segmentations of the LGN. We demonstrated that relying on a single metric, such as the Dice coefficient, provides an incomplete assessment of LGN segmentation quality. By employing a multi-metric framework encompassing quantifying contour, volume, and location, we were able to more accurately characterize the performance of seven current segmentation approaches. Our results indicate that while manual segmentation remains highly accurate, HIPS-THOMAS and DeepThalamus offer the most robust automated alternatives, resulting in high quality LGN segmentations, for high-resolution 7T T1-weighted images. While alignment-based methods like BigBrain can provide reasonable volumetric estimates in a healthy population, they lack the precision required for contour and location accuracy and possibly cannot be applied in a clinical setting. Ultimately, the choice of segmentation method should be guided by the specific requirements of the research application, whether the priority is volumetric estimation, getting a correct contour, or a precise location for the anatomical segmentation.

## Acronyms

AI: Asymmetry Index
baHD: Balanced-Average Hausdorff Distance.
Dice: Dice Coefficient.
HD: Hausdorff Distance.
HIPS-THOMAS: histogram-based polynomial synthesis thalamus optimized multi atlas segmentation
LGN: lateral geniculate nucleus.

## Data availability

Data and custom code that support the findings of this study are available from the corresponding author upon request.

## Author Contributions (CRediT)

**Jan-Luca Schoder**: Data curation, Formal analysis, Methodology, Software, Visualisation, Writing -original draft. **Omer Faruk gulban**: Methodology, Visualisation, Writing -review & editing. **Christa Muller-Axt**: Data preparation, Writing -review & editing. **Rainer Goebel**: Funding acquisition, Methodology, Supervision, Writing -review & editing. **Jan W.Kurzawski**: Data curation, Methodology, Supervision, Visualisation, Writing -review & editing.

