## Supplementary Material for "A critical assessment of LGN segmentation quality based on volume, contour, and location"

### Additional materials

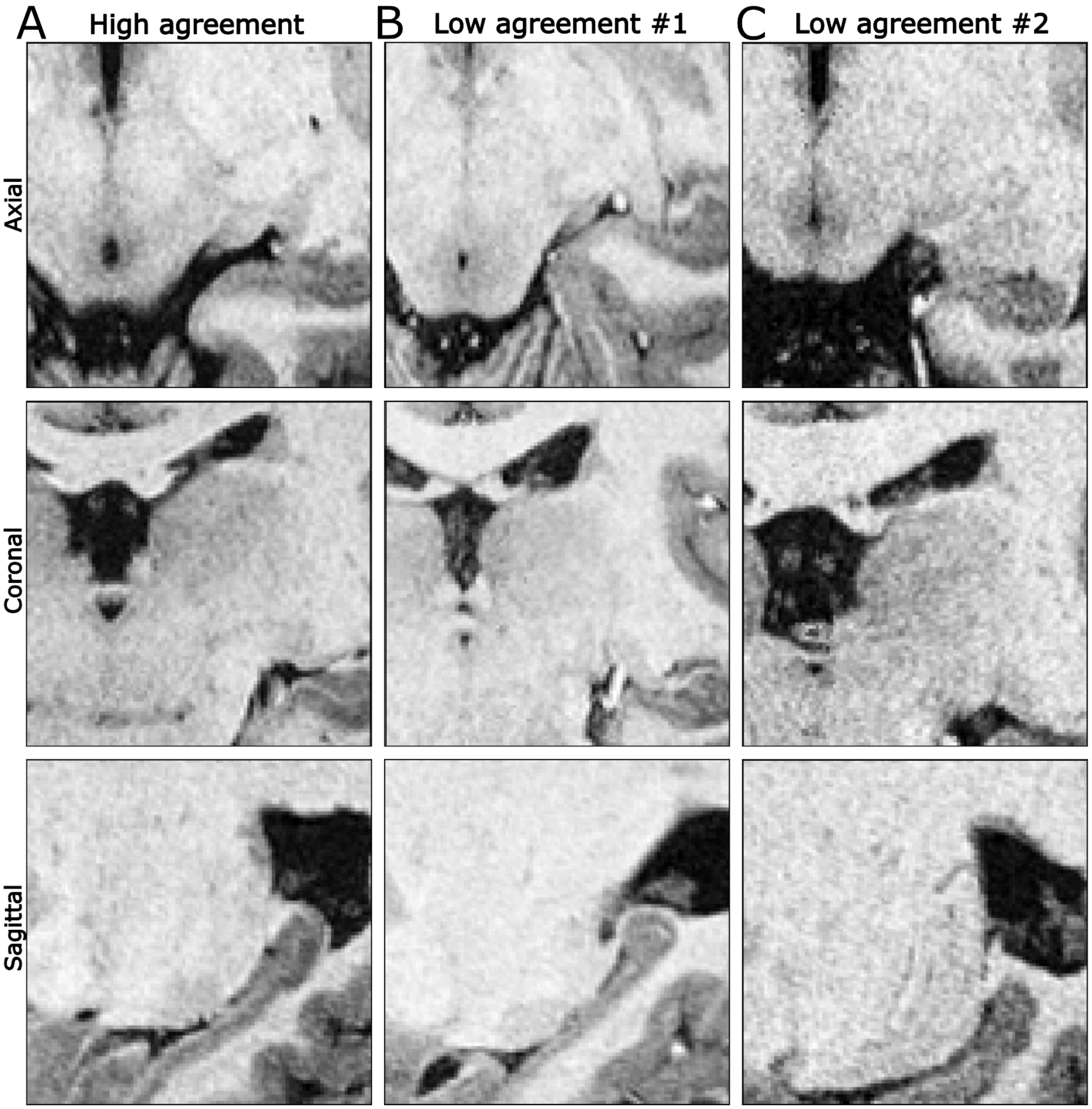


**Fig. S1** *Original T1-weighted images for the overview of highest and lowest segmentation agreement between different segmentation methods.* **A, B** and **C** display an axial, coronal and sagittal slice of example subjects, in correspondence to Fig. 3.

A

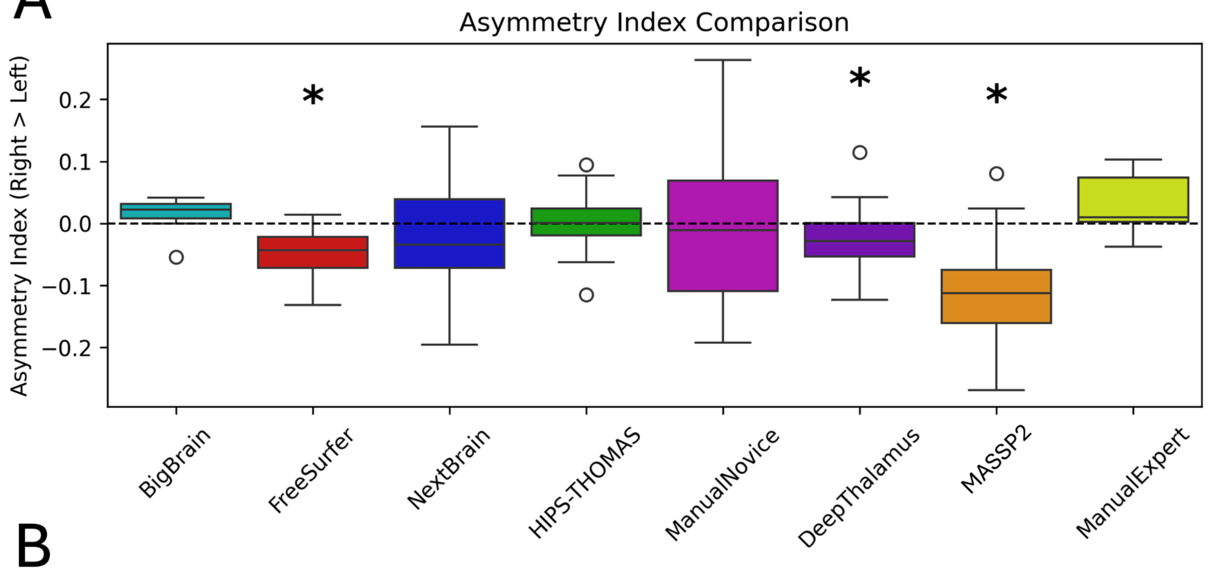


B

| Method | BigBrain | FreeSurfer | NextBrain | HIPS-THOMAS | ManualNovice | DeepThalamus | MASSP2 | ManualExpert |
|---|---|---|---|---|---|---|---|---|
| Median AI | 0.0226 | -0.0438 | -0.0347 | 0.0006 | -0.0111 | -0.0282 | -0.1124 | 0.0094 |
| Q1 (25th percentile) | 0.008 | -0.0715 | -0.0721 | -0.0189 | -0.1093 | -0.0531 | -0.1609 | 0.0027 |
| Q3 (75th percentile) | 0.0317 | -0.0216 | 0.0388 | 0.0238 | 0.0689 | 0.0006 | -0.0748 | 0.0736 |

C

| Method | BigBrain | FreeSurfer | NextBrain | HIPS-THOMAS | ManualNovice | DeepThalamus | MASSP2 |
|---|---|---|---|---|---|---|---|
| Wilcoxon signed rank test | p = 0.1111 | p = 0.00001 | p = 0.0299 | p = 0.0319 | p = 0.0859 | p = 0.0001 | p = 0.00001 |

**Fig. S2** *Asymmetry index of different segmentation methods.* **A:** Boxplot, where boxes represent the median value and the respective Interquartile range. '*' indicates significant (p < 0.0083 (Bonferroni corrected)) difference compared to the manual expert (ground truth) **B**: Table displaying median and the first and third quartile. **C**: Results of the Wilcoxon signed rank test regarding the AI of the segmentation methods compared to the ground truth (Bonferroni-corrected alpha: 0.0083)
